# Exploring Multi-Scale Local and Global Features in Whole Slide Images Using State Space Models

**DOI:** 10.64898/2026.01.26.701877

**Authors:** Chongcong Jiang, Zhuo Zhao, Peixian Liang, Min Shi, Jun Han, Nian-Feng Tzeng, Guanghua Xiao, Danny Z. Chen, Hao Zheng

## Abstract

Whole slide image (WSI) classification is crucial in computational pathology, yet the gigapixel scale of WSIs makes it challenging to extract discriminative and compact WSI-level features for disease diagnosis. In this paper, we propose **MambaWSI**, a novel method that leverages the state space model (SSM) for WSI classification by exploring multi-scale local and global features. Unlike existing approaches that sequentially traverse WSI tiles and rely on vanilla SSMs for long-range dependency modeling, we exploit a traversal strategy in a higher-dimensional discrete space that preserves spatial proximity, enabling a first-local-then-global feature extraction process. Furthermore, to align with the clinical workflow of pathologists when examining WSIs at multiple scales, we propose a two-stage hierarchical fusion strategy: inter-scale feature alignment and aggregation, followed by attention-based fusion across magnifications, integrating complementary information from multiple magnifications. Experiments on two datasets demonstrate that MambaWSI outperforms state-of-the-art methods in classification performance.^1^

## 1 Introduction

Whole slide images (WSIs) contain rich cellular details and micro-environmental context essential for disease diagnosis and prognosis (Cheng et al., 2021; Wang et al., 2024). However, their gigapixel scale poses major computational challenges, and image-level supervision makes it difficult to obtain compact and discriminative WSI representations. Deep learning (DL)-based multiple instance learning (MIL) frameworks have achieved strong performance by dividing WSIs (e.g., 100, 000^2^ pixels) into patches (e.g., 224^2^ pixels each) and aggregating patch-level features for slide-level classification (Gadermayr & Tschuchnig, 2024; Ilse et al., 2018). Various architectures, including convolutional neural networks (CNNs) (Das et al., 2018; B. Li et al., 2021; Lu et al., 2021), graph neural networks (GNNs) (Chen et al., 2021; R. Li et al., 2018), Transformers (C. Jin et al., 2024; Shao et al., 2021), state space models (SSMs) (Fillioux et al., 2023; Yang et al., 2024), and hybrid models (Liang et al., 2024; Liu et al., 2024), have been developed to extract local patch-level features, model region- or WSI-level relationships, and integrate multi-scale information. At the same time, large-scale datasets and self-supervised learning have been leveraged to train pathology foundation models (FMs) (Ahmed et al., 2024; Chen et al., 2024; Huang et al., 2023; Wang et al., 2024). These models demonstrated a strong capability to extract patch-level representations for downstream tasks.

Despite these advances, three challenges remain. **First**, patch-wise partitioning may disrupt spatial relationships, hindering effective modeling of both local and long-range dependencies. GNNs capture local patch interactions but struggle with global context modeling (Chen et al., 2021), while Transformers can model inter-patch correlations but incur quadratic computational costs (Shao et al., 2021). SSMs provide efficient long-sequence modeling, but typically rely on fixed horizontal or vertical traversal to process a WSI, losing spatial coherence (Yang et al., 2024). **Second**, WSIs contain multiple magnification levels, each offering distinct diagnostic insights. But, existing methods lack efficient alignment and fusion strategies to integrate multi-scale information in WSIs while maintaining cross-scale consistency. **Third**, WSIs comprise a large number of redundant patches; attaining a compact yet discriminative WSI-level representation remains challenging. Some methods mitigate this by selecting random or cluster-representative patches to reduce the sample size (Das et al., 2018; Liang et al., 2024; Shao et al., 2021), while others design feature fusion strategies to aggregate information of all patches (Yang et al., 2024). Enhancing the visibility of the most representative instances is crucial to improving WSI classification performance.

To address these challenges, we propose **MambaWSI**, a novel framework that explores multiscale local and global features to derive a compact and discriminative WSI-level representation. **(1)** We leverage Mamba (Gu & Dao, 2023), a variant of SSMs (Gu et al., 2022), to balance computational efficiency and long-sequence modeling. Unlike existing methods, a Z-shaped traversal scheme is applied to non-background patches to ensure a spatially coherent order that first captures local features and then progressively aggregates global context. It enhances locality in feature extraction while preserving long-range dependencies among scattered non-background patches in each WSI. **(2)** Inspired by pathologists’ diagnostic practice, we integrate multiscale features while addressing the inherent mismatch in effective patch numbering across magnification levels. For example, a patch at 5× magnification corresponds to four patches at 10× magnification and 16 patches at 20× magnification. To align features across scales, we employ spatial alignment and random patch discarding, ensuring computational efficiency while reducing overfitting risks. **(3)** We design an attention-based fusion mechanism to aggregate multiscale features for strengthening global contextual relationships while prioritizing highly informative patches, thus obtaining a refined WSI-level representation for final classification. Experiments on two public datasets demonstrate the superior performance of MambaWSI and validate the effectiveness of combining spatially coherent traversal with hierarchical multiscale fusion.

## 2 Methodology

### 2.1 Preliminaries

#### State Space Models (SSMs)

An SSM maps an input state *x*(*t*) ∈ ℝ^*L*^ to an output *y*(*t*) ∈ ℝ^*L*^ via an implicit latent state *h*(*t*) ∈ ℝ^*N*^ . Structured state space sequence (S4) models (Gu et al., 2022) integrate SSMs into DL models using four parameters (**A, B, C, Δ**). The sequence-to-sequence transformation can be represented as a linear ordinary differential equation:

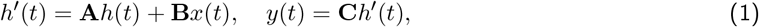

where **A** ∈ ℝ^*N×N*^ is a state transition matrix, **B** ∈ ℝ^*N×*1^ and **C** ∈ ℝ^1*×N*^ are projection matrices, **Δ** is the sampling step size, and *N* is the state size. The learnable continuous parameters **A, B**, and **C** can be discretized by (Gu & Dao, 2023; Gu et al., 2022):

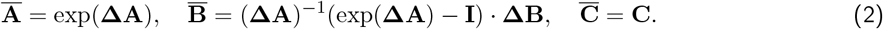

Eq. (1) can thus be re-written in the following recurrent form:

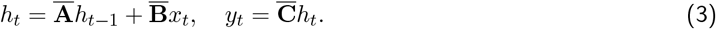

Finally, the convolution can be computed efficiently in parallel:

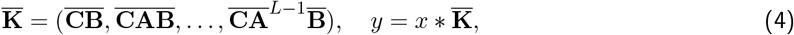

where 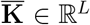 is a structured convolutional kernel and *L* is the sequence length. Mamba (Gu & Dao, 2023) extends the SSM and introduces time-variant parametrization by making **B, C**, and **Δ** dependent on the input, achieving linear complexity with respect to sequence length and effective sequence modeling.

#### Spatial-Filling Curves

Space-filling curves (e.g., Hilbert (Hilbert & Hilbert, 1935) and Z-order (Morton, 1966)) traverse a set of points in a higher-dimensional discrete space without repetition while preserving spatial locality, defined as a bijective function: *ϕ*: ℕ → ℤ^*N*^ , *N* ∈ ℕ. In this paper, MambaWSI leverages the 2D Z-order curve (i.e., *N* = 2), ensuring that nearby patches in the ℤ space remain close after transformation to ℤ^2^. While prior work (Erkan & Aksoy, 2023) used space-filling curves to order WSI patches for position encoding in Transformer models, we integrate Mamba to capture spatial relationships across multiple scales, enabling extraction and fusion of both local and global features.

### 2.2 MambaWSI Overview

Fig. 1 shows an overview of our MambaWSI framework. Given a WSI *X*_*i*_ ∈ ℝ^*H×W*^ and its class label *Y*_*i*_ ∈ {1, 2, … , *C*}, where *C* represents the number of normal/abnormal cases or disease types, we first segment the foreground regions, dividing them into non-overlapping patches at multiple scales. For instance, at 5× magnification, the patches are denoted as 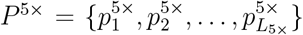, where *L*_5*×*_ is the number of such patches. For simplicity, we omit the magnification notation when the operations apply uniformly across scales. Next, we extract patch-level features using pre-trained networks, such as ResNet (He et al., 2016) pre-trained on ImageNet or pathology-specific FMs (Chen et al., 2024; Huang et al., 2023; Lu et al., 2024). This maps patches into feature representations 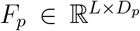 , where *D*_*p*_ is the feature dimension. The extracted features *F*_*p*_ are then fed into the Z-Mamba block, which models spatial relationships across local and long-range dependencies, producing 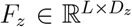 . Finally, features of different scales (i.e., 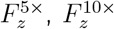, and 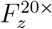) are fused through a hierarchical fusion module to aggregate multi-scale information.

**Figure 1:**
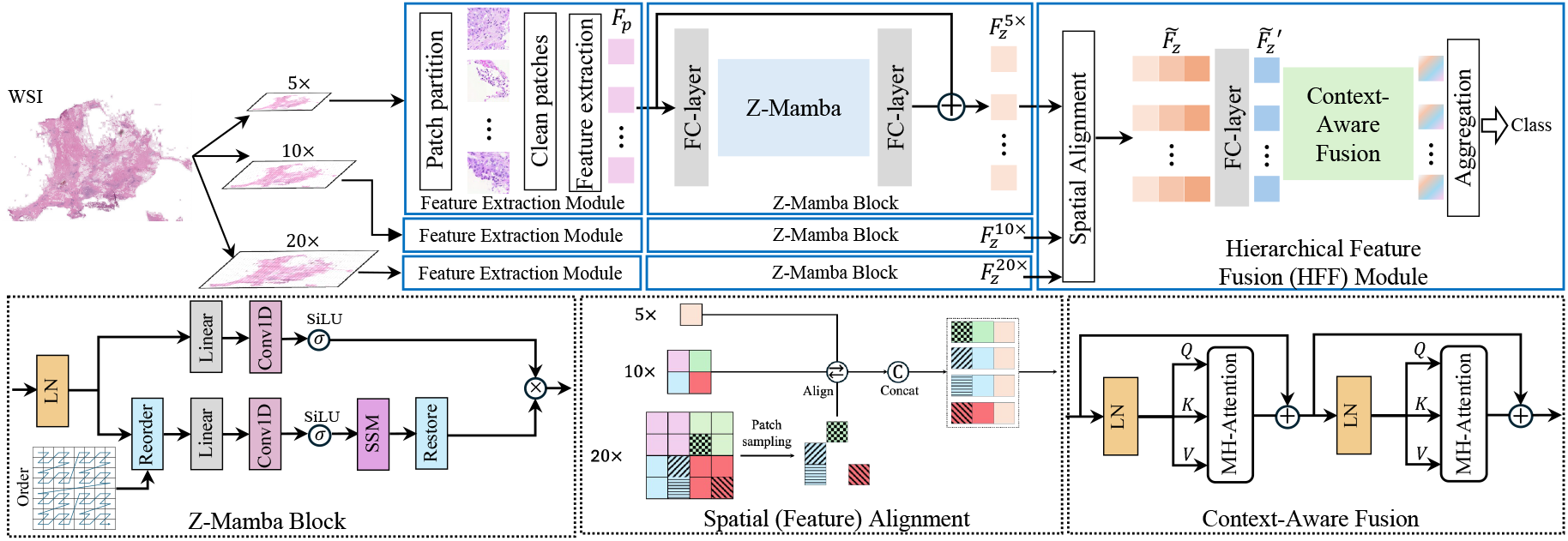
The proposed MambaWSI framework comprises three main steps: (i) Extracting patch-level features from a WSI at three magnifications using a pre-trained feature extractor; (ii) processing multi-scale features in parallel by Z-Mamba blocks to capture both short- and long-range dependencies; (iii) aggregating multi-scale features via a hierarchical fusion module for classification.

### 2.3 Z-Mamba Block: A Locality-Enhanced State Space Module

Unlike Transformers employing positional encoding with input patch sequences, the vanilla Mamba block processes only patch features. This limits the spatial receptive fields and hinders representation learning for non-sequential visual data. Prior approaches attempted to mitigate this issue by applying Mamba in both forward and backward directions (Zhu et al., 2022) or along orthogonal axes (i.e., vertical and horizontal) (Yang et al., 2024), but these strategies did not fully resolve the issue. For example, in a standard row-wise traversal sequence, the first patch in the 2nd row follows the last patch in the 1st row which is geometrically distant from the first patch of the 2nd row. This discontinuity is further exacerbated in large-scale WSIs. Inspired by recent progress (X. Jin et al., 2025), we leverage Z-order traversal to enhance spatial continuity in WSI processing. Consequently, our Z-Mamba block can maintain efficiency in long-sequence modeling while improving spatial feature learning.

For a WSI *X* = *X*_*i*_ ∈ ℝ^*H×W*^ , we divide *X* into *m* × *n* patches of size *R*^2^ each (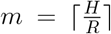and 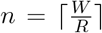), and construct a matrix *S*_*X*_ ∈ ℝ^*m×n*^ that stores the Z-order index *Z*(*a, b*) for each patch at position (*a, b*). *Z*(*a, b*) is obtained by interleaving the binary representations of the row and column indices of each position, forming a unique binary identifier. To retain spatial continuity, we reorder the elements in *S*_*X*_ by sorting them based on the Z-order indices *Z*, yielding an order sequence 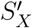 for *S*_*X*_. Further, we remove the background regions to avoid unnecessary computation, resulting in a cleaned sequence 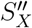 (i.e., 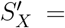 Sort(*S*_*X*_, *Z*), 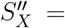 Clean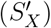). A visualization of 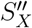 is given in Fig. 2. Patch-level features for non-background regions, 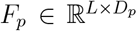 , are extracted by a pre-trained feature extractor and processed by the Mamba module for sequence modeling. Each Mamba block consists of layer normalization (LN) (Ba et al., 2016), convolution, SiLU activation, Selective SSM (Gu & Dao, 2023), and residual connections (He et al., 2016) (see Fig. 1), as:

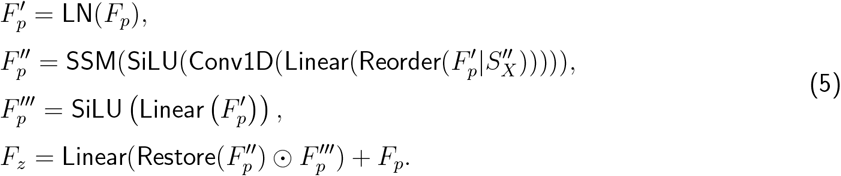

**Figure 2:**
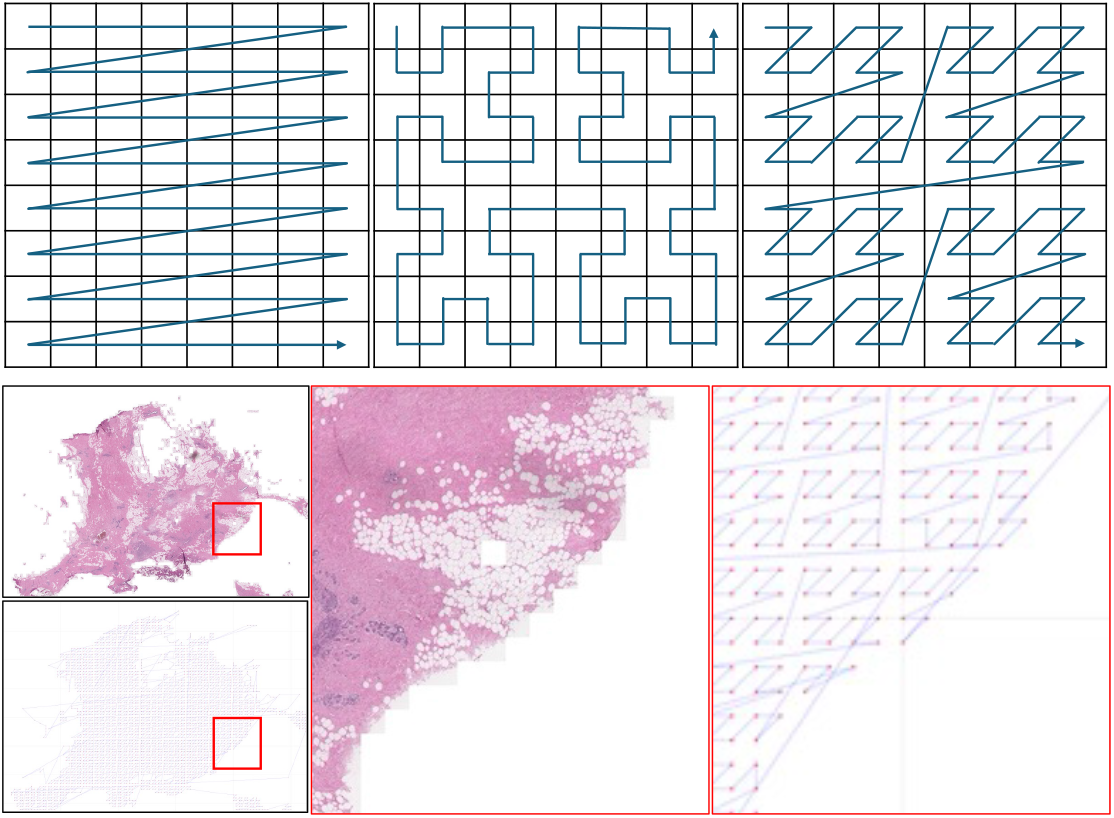
Top: Horizontal, Hilbert, and Z-order scannings. Bottom: foreground patches, a Z-order sequence of patches, and examples.

### 2.4 Hierarchical Feature Fusion Module

The Z-Mamba module processes multi-scale WSI patches in parallel, extracting latent features at different magnifications: 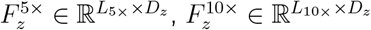 , and 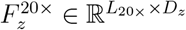 . A straightforward method is to concatenate these features and fuse them using fully connected (FC)-layers, but this method presents three key challenges. (1) Information Imbalance: Since *L*_10*×*_ ≈ 4*L*_5*×*_ and *L*_20*×*_ ≈ 4*L*_10*×*_, finer-scale features may introduce excessive redundancy, increasing computational costs. (2) Spatial Misalignment: The positional correspondence between 5×, 10×, and 20× is neglected. Directly replicating lower-resolution features to match higher-resolution ones also leads to significant computational overhead. (3) Limited Context Awareness: FC-layers struggle to effectively capture inter-patch dependencies.

To address these challenges, we propose a hierarchical feature fusion (HFF) module, which integrates multi-scale features through two progressive stages (see Fig. 1). (1) Spatial (Feature) Alignment: High-resolution features 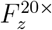 undergo stochastic pruning with a predefined ratio *r*_*d*_, eliminating redundant elements and reducing computational costs (i.e., *L*^*′*^ = (1 − *r*_*d*_)*L*_20*×*_). The retained features, 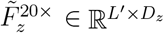, are positionally aligned with their low-resolution counterparts and concatenated channel-wise, producing a unified representation 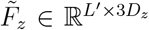 . (2) Context-Aware Fusion: An FC-layer reduces 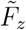 to 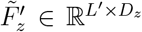 , which is processed by two stacked multi-head attention (MHA) layers with residual connections. The attention mechanism is defined as: 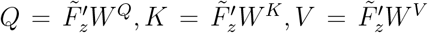 , where 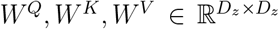 are learnable parameters. For each head *j*, the attention is computed as: 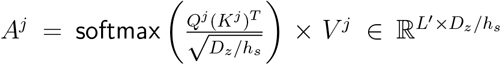, where 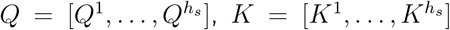, and 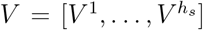, with 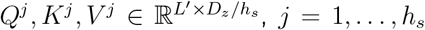, and *h*_*s*_ being the number of heads. Finally, a class token is extracted from 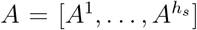 and passed through a linear layer to predict the class probability *q*_*i*_ ∈ [0, 1] for each input sample *X*_*i*_. The model is optimized with the binary cross-entropy loss, as:

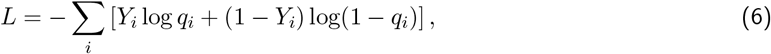

computed over the entire dataset {(*X*_*i*_, *Y*_*i*_)}.

## 3 Experiments

### 3.1 Experimental Setups

We evaluated **MambaWSI** on two public datasets and demonstrated its performance in lung cancer classification. We also conducted ablation studies to assess the effectiveness of its key components.

#### Datasets. (1) TCGA-Lung

A public dataset from the National Cancer Institute Data Portal (Weinstein et al., 2013), containing 1,042 diagnostic WSIs of Lung Squamous Cell Carcinoma (LUSC) and Lung Adenocarcinoma (LUAD) (512 LUSC, 530 LUAD). **(2) NLST-Lung**: A dataset from the National Lung Screening Trial (NLST, 2024), with 572 diagnostic WSIs (228 LUSC, 344 LUAD). All experiments were conducted using five-fold cross-validation (80% for training and 20% for testing in each round). Reported results were averaged over three runs.

#### Evaluation Metrics

We used standard WSI classification evaluation metrics: accuracy (Wikipedia, 2024a) and area under the curve (AUC) (Wikipedia, 2024b).

#### Implementation Details

Our framework was implemented in PyTorch (Paszke et al., 2019) and trained on an NVIDIA A6000 GPU (48 GB). Each WSI was divided into 224^2^ patches at 20×, 10×, and 5× magnifications. The Adam optimizer (Kingma & Ba, 2015) was used with a learning rate of 1*e*^−4^. The optimization stops if the loss saturates after 50 epochs.

#### Comparison Methods

We compared our MambaWSI with state-of-the-art (SOTA) WSI classification methods. ABMIL (Ilse et al., 2018) is an MIL model that aggregates instance features with an attention mechanism to learn instance weights for bag-level prediction. CLAM (Lu et al., 2021) is an attention-based framework that adds a clustering constraint to form class-specific sub-bags and improve separability. DSMIL (B. Li et al., 2021) is a model that processes patches in multiple scales using a dual-stream structure with a consistency loss that aligns instance-level and bag-level predictions. Trans-MIL (Shao et al., 2021) is a Transformer-based model that integrates patch features using self-attention. S4MIL (Fillioux et al., 2023) is the first MIL framework leveraging state space models for WSI classification. MambaMIL (Yang et al., 2024) is an SSM-based approach that processes WSIs using two parallel Mamba branches with horizontal and vertical scannings, followed by fusion via linear layers.

### 3.2 Main Results and Analysis

#### Quantitative Results

Table 1 reports the classification performance of various methods on the two public datasets, using ResNet-50 (He et al., 2016) and PLIP (Huang et al., 2023) as feature extractors. **(I) Comparison with SOTA Methods**. On TCGA-Lung, MambaWSI consistently outperforms all the baseline methods with both feature extractors. With ResNet-50, MambaWSI achieves the highest accuracy 89.2% and AUC 95.1%; with PLIP, it further improves accuracy to 91.5% and AUC to 97.1%. On NLST-Lung, the results indicate a consistent performance advantage of MambaWSI over existing methods, mirroring the trend observed on TCGA-Lung. With ResNet-50, MambaWSI surpasses MambaMIL with a considerable improvement of 2.1% in accuracy and 2.3% in AUC, reinforcing its robustness across the datasets. **(II) Comparison across Feature Extractors**. ResNet-50 was pre-trained on generic images, while PLIP is a pathology-specific foundation model. Across all the methods, PLIP-extracted features led to consistently superior results compared to ResNet-50, highlighting the advantage of domain-specific pre-training for WSI classification. Further, despite the feature extractor choices, MambaWSI consistently outperforms alternative methods, demonstrating that a well-designed framework for integrating instance features into WSI-level representations is crucial to higher performance. Overall, these results highlight MambaWSI’s superiority in WSI classification, demonstrating its better ability to capture spatial dependencies and multi-scale features than SOTA MIL frameworks based on CNNs (Ilse et al., 2018), Transformers (Shao et al., 2021), or SSMs (Fillioux et al., 2023; Yang et al., 2024).

**Table 1:** Performance comparison of different methods using two feature extractors on the TCGA-Lung and NLST-Lung datasets.

|  | Dataset<br>Method | TCGA-Lung |  | NLST-Lung |  |
| --- | --- | --- | --- | --- | --- |
| | | Accuracy ↑ | AUC ↑ | Accuracy ↑ | AUC ↑ |
| ResNet-50 | Mean-Pooling | 0.853 ± 0.021 | 0.925 ± 0.015 | 0.764 ± 0.041 | 0.840 ± 0.043 |
| | Max-Pooling | 0.875 ± 0.021 | 0.944 ± 0.017 | 0.885 ± 0.027 | 0.940 ± 0.031 |
| | ABMIL (Ilse et al., 2018) | 0.873 ± 0.015 | 0.935 ± 0.017 | 0.872 ± 0.021 | 0.923 ± 0.015 |
| | CLAM (Lu et al., 2021) | 0.872 ± 0.018 | 0.937 ± 0.014 | 0.860 ± 0.025 | 0.937 ± 0.017 |
| | DSMIL (B. Li et al., 2021) | 0.879 ± 0.024 | 0.945 ± 0.020 | 0.880 ± 0.025 | 0.943 ± 0.017 |
| | TransMIL (Shao et al., 2021) | 0.876 ± 0.021 | 0.944 ± 0.017 | 0.874 ± 0.032 | 0.914 ± 0.028 |
| | S4MIL (Fillioux et al., 2023) | 0.870 ± 0.019 | 0.937 ± 0.015 | 0.879 ± 0.019 | 0.940 ± 0.016 |
| | MambaMIL (Yang et al., 2024) | 0.884 ± 0.023 | 0.946 ± 0.016 | 0.894 ± 0.015 | 0.942 ± 0.023 |
|  | **MambaWSI (Ours)** | **0.892 ± 0.010** | **0.951 ± 0.014** | **0.915 ± 0.021** | **0.965 ± 0.026** |
| PLIP | Mean-Pooling | 0.886 ± 0.025 | 0.949 ± 0.022 | 0.840 ± 0.026 | 0.910 ± 0.011 |
| | Max-Pooling | 0.878 ± 0.021 | 0.950 ± 0.016 | 0.901 ± 0.024 | 0.964 ± 0.026 |
| | ABMIL (Ilse et al., 2018) | 0.901 ± 0.029 | 0.964 ± 0.019 | 0.920 ± 0.018 | 0.973 ± 0.015 |
| | CLAM (Lu et al., 2021) | 0.907 ± 0.022 | 0.964 ± 0.016 | 0.907 ± 0.021 | 0.972 ± 0.015 |
| | DSMIL (B. Li et al., 2021) | 0.905 ± 0.024 | 0.965 ± 0.015 | 0.904 ± 0.024 | 0.971 ± 0.014 |
| | TransMIL (Shao et al., 2021) | 0.891 ± 0.027 | 0.958 ± 0.016 | 0.916 ± 0.029 | 0.970 ± 0.020 |
| | S4MIL (Fillioux et al., 2023) | 0.892 ± 0.020 | 0.964 ± 0.018 | 0.910 ± 0.021 | 0.970 ± 0.015 |
| | MambaMIL (Yang et al., 2024) | 0.907 ± 0.023 | 0.967 ± 0.014 | 0.914 ± 0.028 | 0.972 ± 0.016 |
|  | **MambaWSI (Ours)** | **0.915 ± 0.017** | **0.971 ± 0.010** | **0.923 ± 0.021** | **0.977 ± 0.008** |

#### Qualitative Results

Fig. 3(a) provides a t-SNE (Van der Maaten & Hinton, 2008) visualization of WSI-level representations obtained by MambaMIL and our MambaWSI model, respectively, highlighting its abilities to enhance intra-class similarity, reduce inter-class overlap, and produce more discriminative representations.

**Figure 3:**
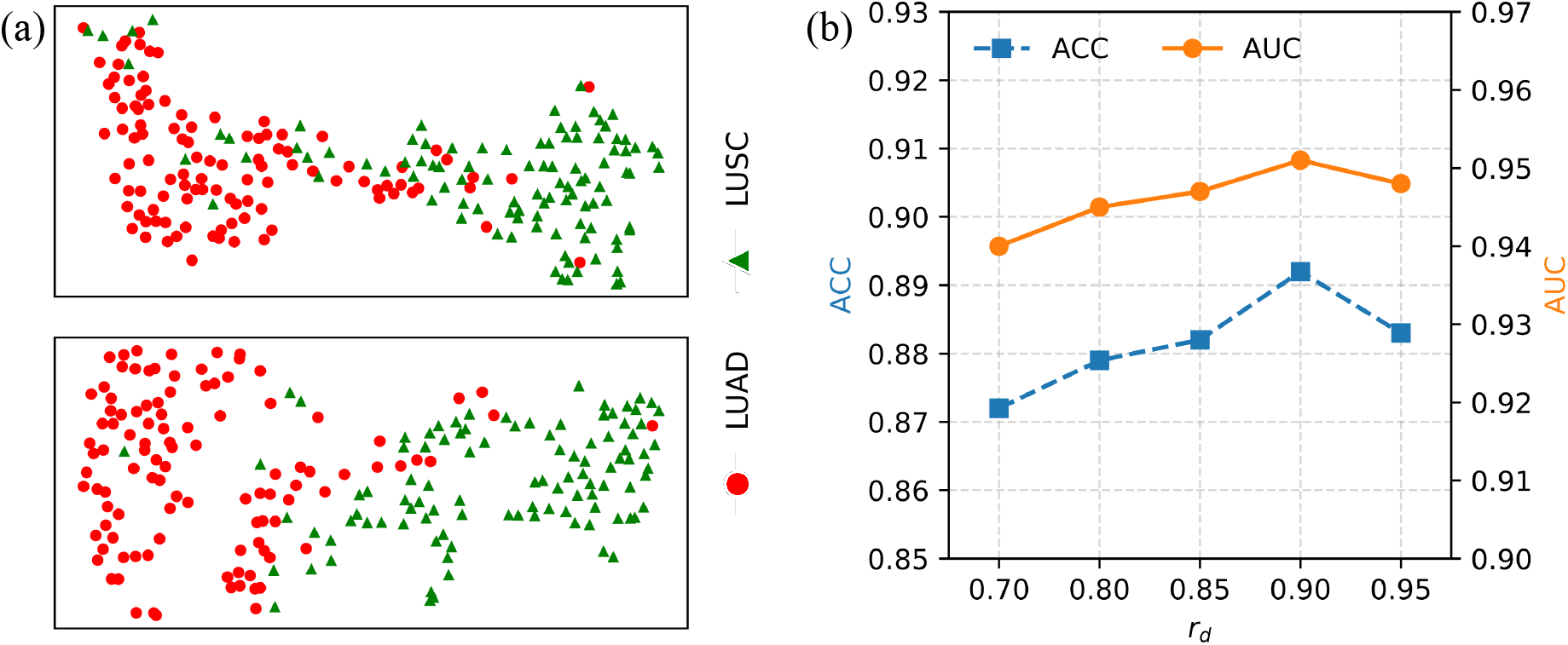
(a) t-SNE visualization of WSI-level representations learned by MambaMIL (top) and MambaWSI (bottom) on the TCGA-Lung dataset. (b) MambaWSI performance with different values of *r*_*d*_.

#### 3.3 Ablation Studies

#### Traversal Orders

We evaluated the impact of patch traversal order in MambaWSI. Fig. 2 illustrates three traversal strategies. Table 2 shows that the Z-order traversal performs the best, suggesting that preserving spatial locality information benefits the Mamba module.

**Table 2:** Ablation studies on NLST-Lung using ResNet features.

| | Setting | Accuracy ↑ | AUC ↑ |
| --- | --- | --- | --- |
| Order | Horizontal | 0.870 ± 0.024 | 0.942 ± 0.016 |
| | Hilbert | 0.893 ± 0.019 | 0.947 ± 0.011 |
| | Z-order | 0.909 ± 0.017 | 0.959 ± 0.010 |
| Scale | 5× | 0.881 ± 0.027 | 0.940 ± 0.019 |
| | 10× | 0.893 ± 0.025 | 0.955 ± 0.017 |
| | 20× | 0.909 ± 0.017 | 0.959 ± 0.010 |
| | 5× & 10× | 0.901 ± 0.029 | 0.958 ± 0.024 |
| Fusion | Concat | 0.881 ± 0.017 | 0.941 ± 0.029 |
| | Replicate | 0.904 ± 0.025 | 0.960 ± 0.031 |
| **Ours** | Z-order<br>5× & 10× & 20×<br>HFF | **0.915** ± 0.021 | **0.965** ± 0.026 |

#### Single vs Multi-scale

We examined the effect of using different scales by removing some scales from our MambaWSI while keeping the FC-layer and MHA blocks intact. As shown in Table 2, among single-scale settings, 20× achieves the best performance, and 10× surpasses 5×. The dual-scale configuration 5× & 10× yields better performance than either single-scale 5× or 10× alone, but still falls short of our three-scale setting. These results highlight the advantage of multi-scale feature aggregation.

#### Fusion Mechanisms

We investigated three strategies for fusing features. (i) Concat: It directly concatenates the feature matrices, yielding features in 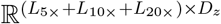. (ii) Replicate: It aligns 5× and 10× features with the 20× grid using hierarchical parent maps, replicates the aligned features, and concatenates them with the 20× features, yielding features in 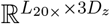, which are then linearly projected to 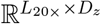. (iii) Our HFF scheme: It produces features in 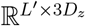 , with *L*^*′*^ ≪ *L*_20*×*_. The results in Table 2 indicate that simple concatenation fails to capture patch correlations, leading to inferior performance. Replication improves accuracy but incurs significant computational overhead. Our HFF scheme yields the best performance while maintaining efficiency.

#### Discarding Ratio *r*_*d*_

Fig. 3(b) shows that we experimented with different ratios (the *x*-axis) and chose *r*_*d*_ = 0.9 in our MambaWSI.

## 4 Conclusions

We proposed MambaWSI, a novel framework for WSI classification that integrates a Z-order traversal strategy and multi-scale feature fusion via the state space module. By feeding Z-ordered patch features into the Mamba module, our MambaWSI preserves spatial proximity while capturing long-range dependencies. Multi-scale feature alignment enhances the local feature representation, and hierarchical fusion efficiently integrates complementary information across multiple magnifications using an attention-based mechanism. Experiments on two public datasets demonstrated the superiority of our MambaWSI over state-of-the-art alternative methods and validated the effectiveness of its key components.

## 5 Compliance with ethical standards

This research was conducted retrospectively using human subject data made available in open access (NLST, 2024; Weinstein et al., 2013). Ethical approval was not required, as confirmed by the license attached to the public data.

## Footnotes

1 This work has been submitted to the IEEE. Copyright may be transferred without notice, after which this version may no longer be accessible.

